# A new inverse probability of selection weighted Cox model to deal with outcome-dependent sampling in survival analysis

**DOI:** 10.1101/2023.02.07.527426

**Authors:** Vera H. Arntzen, Marta Fiocco, Inge M.M. Lakeman, Maartje Nielsen, Mar Rodríguez-Girondo

**Affiliations:** Mathematical Institute, Leiden University, Leiden, The Netherlands; Department of Medical Statistics and Bioinformatics, Leiden University Medical Center, Leiden, The Netherlands; Department of Clinical Genetics, Leiden University Medical Center, Leiden, The Netherlands; Department of Human Genetics, Leiden University Medical Center, Leiden, The Netherlands

**Keywords:** survival analysis, outcome-dependent sampling, weighting, Cox regression, genetic epidemiology

## Abstract

Motivated by the study of genetic effect modifiers of cancer, we examined weighting approaches to correct for ascertainment bias in survival analysis. Family-based outcome-dependent sampling is common in genetic epidemiology leading to study samples with too many events in comparison to the population and an overrepresentation of young, affected subjects. A usual approach to correct for ascertainment bias in this setting is to use an inverse probability-weighted Cox model, using weights based on external available population-based age-specific incidence rates of the type of cancer under investigation. However, the current approach is not general enough leading to invalid weights in relevant practical settings if oversampling of cases is not observed in all age groups. Based on the same principle of weighting observations by their inverse probability of selection, we propose a new, more general approach. We show the advantage of our new method using simulations and two real datasets. In both applications the goal is to assess the association between common susceptibility loci identified in Genome Wide Association Studies (GWAS) and cancer (colorectal and breast) using data collected through genetic testing in clinical genetics centers.

## 1 INTRODUCTION

Family-based outcome-dependent sampling is common in genetic epidemiology. Since harmful variants in cancer associated high risk genes are typically rare, an efficient sampling strategy to find carriers of these variants is to oversample affected individuals with a family history of a specific disease. For example, carriers of pathogenic variants in the Lynch syndrome associated gene *PMS2* and the breastand ovarian cancer associated genes *BRCA1* and *BRCA2*, are often detected through genetic screening programs in which testing is targeted to families with multiple cases. Due to this testing strategy, the available study cohorts to investigate modifiers of cancer risk are often non-representative samples of the population of interest: families with members that are diagnosed at a young age, and multiple cases are more likely to be included in the sample.

In the context of survival analysis, family-based outcome-dependent sampling results in an overrepresentation of events and short lifetimes, which without adjustment, leads to biased estimates of covariate effects when using, for example, a Cox proportional hazards model. This happens because the sampling mechanism affects the joint distribution of the time-to-event and covariate.

To solve this problem, two main approaches have been proposed in the literature: methods based on retrospective likelihood (1–3) and the weighted cohort method (4) based on weighted Cox regression. The general idea of the methods based on retrospective likelihood is to formulate the likelihood of the observed covariate values conditional on the observed outcomes. These methods typically require to know the familial relations within the sample and the distribution of the covariate of interest, leading to analytically complex and computationally intensive methods. When the overall age-specific incidence rates in the population of interest are known, an alternative approach to estimate the association between a set of covariates and time to cancer diagnosis under outcome-dependent sampling is to use a weighted Cox regression model (4). The general idea is to propose a weighting scheme with different weights for affected (observed events) and unaffected (right-censored) individuals according to an external source, so that the resulting weighted sample mimics the true target population (4, 5) in terms of the age-specific proportions of affected and unaffected individuals. Due to its simplicity, this is an attractive approach. However, the proposed weighted scheme has some limitations: it often leads to invalid weights in relevant practical situations since it is only workable under particular sampling schemes, for example with strong oversampling of cases only.

This study has two objectives. Firstly, we propose an alternative, more general, inverse probability of selection weighting scheme using population-based age-specific incidence rates of the event of interest. Accordingly, a generalized weighted cohort method that can deal with arbitrary levels of outcome-dependent sampling is proposed and compared with the existing approach. A second goal of this study is to investigate the role of within-family correlation due to unobserved shared factors in the performance of weighted cohort approaches. Despite the presence of multiple members of the same family being relatively common in studies relying on the original weighted cohort method (see Table S1 for details), the impact of unobserved heterogeneity in this context has not been previously evaluated. It is an aspect that deserves attention.

The rest of the paper is organised as follows. In Section 2, the commonly used weighted cohort Cox approach is revisited and its assumptions are discussed. A new alternative weighting scheme is proposed in Section 3. In Section 4, both weighting schemes are compared by means of an intensive Simulation study. In Section 5, we present two real data illustrations. In both illustrations, the role of genetic variants as modifiers of cancer risk is studied using datasets of affected individuals and family members ascertained through genetic counseling in a clinical genetic center. In the first application, we focus on colorectal cancer in carriers of the pathogenic variant *PMS2*, and in the second application, we analyse the association between a Polygenic

Risk Score (PRS) based on common breast cancer-associated variants and breast cancer risk in multiple case families. Main conclusions, recommendations, and a final discussion follow in Section 6.

## 2 WEIGHTED COX REGRESSION TO DEAL WITH OUTCOME-DEPENDENT SAMPLING

Let *T* be the time to event of interest in the target population of interest and denote by *C* the right censoring time, assumed to be uninformative. Denote by *Z* the covariate of interest. Since sampling schemes in genetic epidemiology are typically family-based, denote the observed sample information by (*t*_*ij*_, *δ*_*ij*_, *z*_*ij*_), where *i* = 1, …, *n*_*j*_ index all included individuals in the sample belonging to family *j*, and assume that *M* families are observed, with varying observed size *n*_1_, …, *n*_*M*_, so that 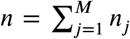 individuals are included in the sample. The observed time to event for individual *ij* is given by *t*_*ij*_ = min(*T*_*ij*_, *C*_*ij*_). Define the non-censoring indicator *δ*_*ij*_ = *I*(*T*_*ij*_ ≤ *C*_*ij*_) where *δ*_*ij*_ is 1 if the event is observed or 0 if observation *ij* is right censored. *z*_*ij*_ denotes covariate value for individual *ij*.

The observed data is collected through an outcome-dependent sampling scheme. The specific selection criteria vary over studies, depending on the severity and prevalence of the disease, but are usually of the form 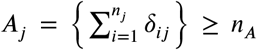. That is, if and only if there are at least *n*_*A*_ family members in family *j* who experienced the event of interest, family *j* is included in the study. Often, not only *δ*_*ij*_, *i* = 1, …, *n*_*j*_ but also *t*_*ij*_, *i* = 1, …, *n*_*j*_ is used to define the ascertainment event *A*_*j*_. For example, including families with at least three members with cancer diagnosis below the age of 60 would correspond to 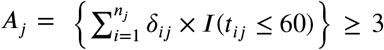. In general, if a family *j* is included in the study, i.e. if *A*_*j*_ = 1, all its members are invited to participate, also those who did not experience the event yet (right-censored observations). Despite the diversity of possible family configurations, there is typically an overrepresentation of young cases of the family. Often, only the carriers of a certain genetic pathogenic variant are of interest and the target population is the population of carriers of such a variant. In those situations, the former definitions still apply, but the family size is restricted to the family members who are mutation carriers.

A common approach to estimate the effect of covariate *Z* on *T* is to use the Cox proportional hazards model with hazard function *h*(*t*|*z*) = *h*_0_(*t*) exp(*βz*) where *h*_0_(*t*) is the baseline hazard. With prospective cohort data, the parameter *β* can be estimated maximizing the partial likelihood. However, the overrepresentation of events and short event times in the sample due to outcome-dependent sampling alters the risk set composition along the follow-up time in comparison to the true population, which may result in biased estimation of the covariate effect. A possible solution to this problem is to consider a weighted Cox model using external information about the distribution of *T* in the population to construct weights reflecting individuals’ selection probabilities.

### 2.1 The weighted cohort approach revisited

When *T* represents the age at cancer diagnosis, or another common disease, registry data about the distribution of *T* in the target population is often available. Let us assume, as it is common in practice, that the available information in the external source is aggregated into *K* disjoint age intervals, defined as *I*_1_ = [*α*_0_, *α*_1_), *I*_2_ = [*α*_1_, *α*_2_), … *I*_*K*_ = [*α*_*K*−1_, *α*_*K*_). In the case of cancer studies, the typical available external information is the population cancer incidence rate *μ*_*k*_ for each age interval *I*_*k*_, *k* = 1, … *K*. The seminal paper of *Antoniou et al. ((***?** )) proposed a weighted Cox regression model with sampling weights derived such, that the incidence rates in each interval *I*_*k*_ in the resulting pseudo-population after weighting agrees with the incidence rates *μ*_*k*_ in the target population, available and used as external data to calculate the weights.

Specifically, let *r*_*k*_ denote the number of individuals experiencing the event within the age interval, *I*_*k*_ = [*α*_*k*−1_, *α*_*k*_), *k* = 1, … *K*. Similarly, *s*_*k*_ denotes the number of individuals right-censored within the age interval *I*_*k*_ (i.e. follow-up ends between age *α*_*k*−1_ and *α*_*k*_ without the event being observed). The term 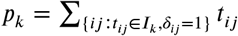 denotes the total follow-up time accumulated by all *r*_*k*_ individuals experiencing the event in age interval *I*_*k*_; the equivalent total follow-up time accumulated by the *s*_*k*_ rightcensored individuals is denoted by 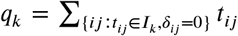. Then, all *r*_*k*_ cases in interval *I*_*k*_ are assigned weight *w*_*k*_ and all *s*_*k*_ right-censored individuals in interval *I*_*k*_ are assigned weight *v*_*k*_ such that:

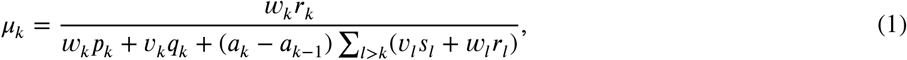

In the right part of expression (1) the weighted total of affected observations is divided by the weighted total of observations at risk. Then, this weighted ratio is imposed to be equal to the population incidence rate *μ*_*k*_. As a result, after weighting the sample age-specific incidence rates resemble the age-specific incidence rates of the population. However, since equation (1) alone does not guarantee unique weights *w*_*k*_ and *v*_*k*_, the following constraint is incorporated to guarantee unique weights:

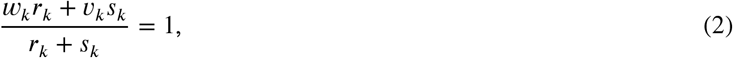

Combining equations (1) and (2) provides unique expressions for *v*_*k*_ and *w*_*k*_:

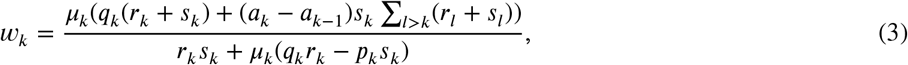

where Σ_*l*>*k*_(*r*_*l*_ + *s*_*l*_) are all observations in age groups older than *k*. The weight equation for censored individuals is given by

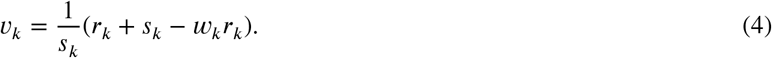

Once weights *v*_*k*_, *w*_*k*_ for each age interval *I*_*k*_, *k* = 1, …, *K* are calculated, the regression parameter *β* can be estimated using the following weighted score equation:

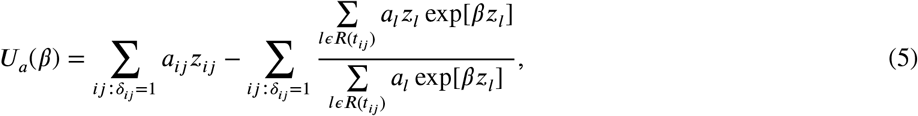

where *R*(*t*_*ij*_) is the set of individuals that are still at risk just before *t*_*ij*_, i.e. *R*(*t*_*ij*_) = {*l* : *t*_*ij*_ ≤ *t*_*l*_}, and weight *α*_*ij*_ for individual *ij* (*i* = 1, …, *n*_*j*_, *j* = 1, …, *M*) is defined as

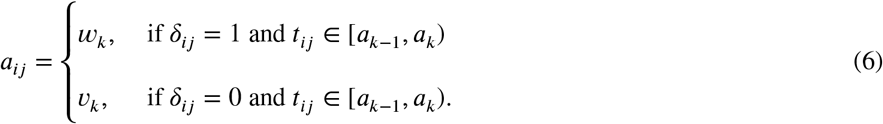

A number of conditions are required to guarantee finite and positive weights *w*_*k*_ and *v*_*k*_, namely:

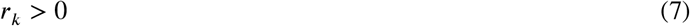

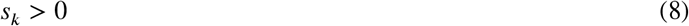

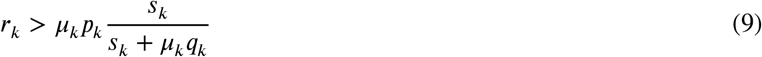

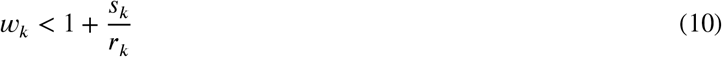

Conditions (7) and (9) are required to get valid *w*_*k*_ weights for the cases, while conditions (8) and (10) are required to get valid *v*_*k*_ weights for those that are censored. Condition (7) implies the observation of events in all the considered intervals and condition (8) implies the presence of right-censored observations in all considered intervals. Conditions (9) and (10) are more difficult to interpret and evaluate beforehand, but they are both related to the level of oversampling of events. As discussed by *Antoniou et al. ((***?** )), if oversampling of events occurs in all considered age groups both conditions are typically fulfilled. However, as we will show in our real data application, oversampling of young cancer cases is the norm in genetic epidemiology, but not necessarily the case at older ages, so condition (9) and especially (10) might not be fulfilled in relevant practical scenarios. Under oversampling of events at interval *I*_*k*_, condition (9) is verified since *r*_*k*_ > *μ*_*k*_*p*_*k*_, i.e., the observed number of events in interval *I*_*k*_ is larger than the expected number of events assuming the population incidence rate (*μ*_*k*_). Actually, since *μ*_*k*_*q*_*k*_ is usually positive, 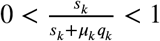 in general which implies that condition (9) is fulfilled even when no oversampling of events is observed in interval *I*_*k*_. However, condition (10) is cumbersome. Since it involves the estimated weight for events *w*_*k*_ together with the ratio of events and right-censored observations in interval 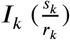, this condition is often not satisfied when there is no clear oversampling of cases in interval *I*_*k*_. In such situations, *w*_*k*_ can still be calculated but it becomes small, which leads to violating condition (10).

When any of the conditions (7)-(10) are not satisfied, the weighted cohort method can still be applied by merging intervals, however the method becomes then less precise and dependent on sample specific characteristics which may hamper comparability among studies using this method.

Next, we propose an alternative, more general weighting scheme which allows to overcome the aforementioned limitations.

### 2.2 The new generalised weighted cohort approach

We propose a new weighting scheme to correct outcome-dependent sampling using external information. In contrast to the previous weighted cohort method, the new approach is more general, as it can be applied with arbitrary levels of over or underrepresentation of events.

Similar to the original method, the newly proposed weights represent sampling probabilities given the observed time to event of each individual so that the resulting pseudo-population matches the target population of reference in terms of the distribution of *T*. However, here we take a different approach to derive the weights. Instead of directly using the incidence rates, we focus on the risk sets at the beginning at each interval *I*_*k*_ and weight the individuals so that the resulting weighted risk set presents the same ratio of events and non-events as one would expect if the sample would have been randomly drawn from the target population.

Let *N*_*k*_ denote the number of individuals at risk (those who did not experience the event yet) at the beginning of the interval *I*_*k*_ in our sample, denoted by *𝒮*_*O*_, potentially drawn under an outcome-dependent sampling mechanism. Now denote by *𝒮*_*P*_ a hypothetical random sample of the target population with the same *N*_*k*_ number of individuals at risk at the beginning of the interval *I*_*k*_. In both cases, *N*_*k*_ can be split into two disjoint parts: a) the number of individuals that experience the event within the interval *I*_*k*_ and b) those experiencing the event in later intervals. However, if *𝒮*_*O*_ is obtained using outcome-dependent sampling, the expected number of individuals belonging to each of these two parts in *𝒮*_*O*_ and *𝒮*_*P*_ will, in general, differ. For the hypothetical random sample *𝒮*_*P*_, *N* can be decomposed as follows:

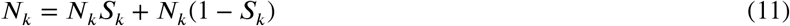

where *S*_*k*_ = *P* (*T* > *α*_*k*_| *T* > *α*_*k*−1_) represents the conditional probability of experiencing the event in a later time interval than interval *I*_*k*_ given that the event has not been experienced before interval *I*_*k*_ in the reference population. *S*_*k*_ can be directly calculated from the typically available population cancer incidence rates *μ*_*k*_ for each age interval *I*_*k*_, since 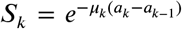, *k* = 1, … *K*. Accordingly, 1 − *S*_*k*_ is the probability of experiencing the event in the interval *I*_*k*_ given that it has not been experienced before, in the reference population. From expression (11) follows that the ratio between events and non-events in interval *I*_*k*_ in the reference population is given by 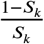.

The same decomposition of the risk set at the beginning of interval *I*_*k*_ can be made for the observed sample *𝒮*_*O*_, potentially subject to outcome-dependent sampling:

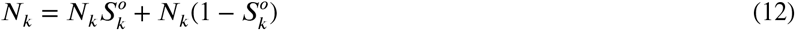

where 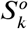 is the observed proportion of individuals at risk at time *α*_*k*−1_ experiencing the event beyond *I*_*k*_, calculated with the sample data.

In our new approach, we keep those subjects non-experiencing the event at interval *I*_*k*_ unweighted (*v*_*k*_ = 1) while we assign specific weights (*w*_*k*_) to those subjects experiencing the event of interest in interval *I*_*k*_ making use of the decompositions given by expressions (11) and (12). Specifically, weights *w*_*k*_ correct the oversampling (or undersampling) of cases, such that the ratio between events and non-events in interval *I*_*k*_ in the resulting pseudo-population after weighting is the same as in the reference population:

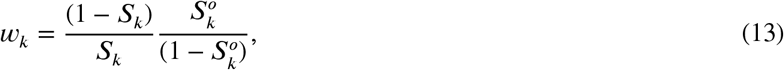

Equation (13) indicates that the population ratio between events and non-events in the interval *I*_*k*_, given by 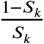 is multiplied by the inverse quantity based on the observed data,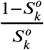. After weighing, the composition of the risk set at interval *I*_k_ resembles the composition of the risk set at interval *I*_*k*_ in the reference population. As a result, under oversampling of cases, weights for affected individuals in interval *I*_*k*_ are *w*_*k*_ < 1, representing the inverse of the probability of being selected. Alternatively, under undersampling of cases, *w*_*k*_ > 1. Interestingly, in absence of outcome-dependent sampling, i.e. under random sampling, *w*_*k*_ = 1 and the new method coincides with the regular unweighted Cox model.

With our new proposal, two conditions need to be fulfilled in order to get valid weights: 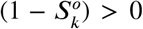 and *S*_*K*_ > 0. The first condition 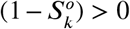 is satisfied if events are observed in each interval *I*_*k*_, so as the original weighted cohort method, observation of events in all group ages is a requirement of our new method. However, the new method does not require the presence of right-censoring which makes it a more general and natural approach. The condition *S*_*K*_ > 0 only involves the last interval and implies that the method is suitable for studying events not experienced by a part of the population during the relevant follow-up time. This is a mild condition that is always satisfied when studying defective distributions (*S*(∞) > 0) such as time to cancer or other diseases since not all population members will develop the event of interest. Even if our interest would be to study time to death or the target population would be a highly susceptible population to a specific cancer with lifetime risk of 1, the new weights could still be applied with an appropriate choice of the upper limit of the last interval *K*.

Once the weights are calculated, the regression parameter *β* can be estimated using the weighted score equation (5).

In summary, both the existing weighted cohort and the new generalised weighted cohort approaches generate pseudo populations by means of inverse probability of selection weighting, but these subpopopulations are different. The new method is more general since it does not require oversampling or undersampling of events in all or at specific intervals and does not make assumptions about the right-censoring distribution.

### 3 SIMULATION STUDY

A simulation study was conducted to assess the new generalized weighted cohort method’s performance and compare it with the existing approach in several scenarios intended to mimic relevant situations in practice. We consider two main simulation settings. First, we generate data under the assumption of fully observed heterogeneity, i.e., differences among individuals in terms of hazards can be fully accounted for by the observed covariates. In a second simulation setting, we consider the presence of unmeasured factors shared within families.

### 3.1 Simulation setup I

Simulated data was generated using the following model:

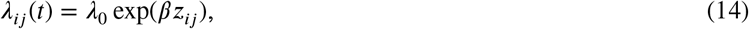

where *t* is the observed event time, 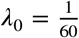 represents the constant baseline hazard, *Z* is a continuous covariate assumed to be normally distributed (*Z* ∼ *N*(0, 1)) and *β* its associated log-hazard ratio. If the resulting event times were larger than 100, these were set to 100. Censoring times were sampled from an exponential distribution (*C* ∼ *exp*(60)) and the family size in the population is set to *n*_*j*_ (family size of size *n*_*j*_ = 2 and 5 members were considered). In each Monte Carlo trial, we generated *M* = 1000 families (*M* = 125, 250, 500). Family-based outcome-dependent sampling was implemented by including families in the sample if for at least *n*_*A*_ family members the event was observed before the end of follow-up (*n*_*A*_ = 1, 3). The different combinations of *n*_*A*_ and *n*_*j*_ lead to three different scenarios with increasing level of outcome-dependent sampling: scenario 1 (A1) with *n*_*j*_ = 5 and *n*_*A*_ = 1 represents the mildest level of selection, scenario 2 (A2) with *n*_*j*_ = 5 and *n*_*A*_ = 3 represents a medium level of outcome dependent selection, and scenario 3 (A3) with *n*_*j*_ = 2 and *n*_*A*_ = 1 represents the strongest level of outcome dependent sampling in the simulation study. Moreover, all included families had at least one ‘young affected’ defined as having observed event time smaller than the first quartile of simulated *T* distribution. This mimics the common practice in clinical genetics centers: families are invited to participate in genetic studies when a young family member is diagnosed with the event at a young age. In terms of covariate effect, the null case (*β* = 0) and two alternative scenarios (*β* = 0.3, 1) were considered.

For each considered value of *β*, the underlying population was generated simulating a large data set (*N* = 200 000) without ascertainment and it was used to approximate the population hazards needed to calculate the weights. Bias, mean square error and coverage proportions of the 95% confidence intervals are reported in Table 1. As proposed by *Antoniou et al. (2005)*, robust estimates of the standard errors were obtained using a sandwich estimator in both considered weighted methods. Moreover, the proportion of invalid weights in the *M* Monte Carlo trials is reported for the two weighted methods.

**Table 1.**
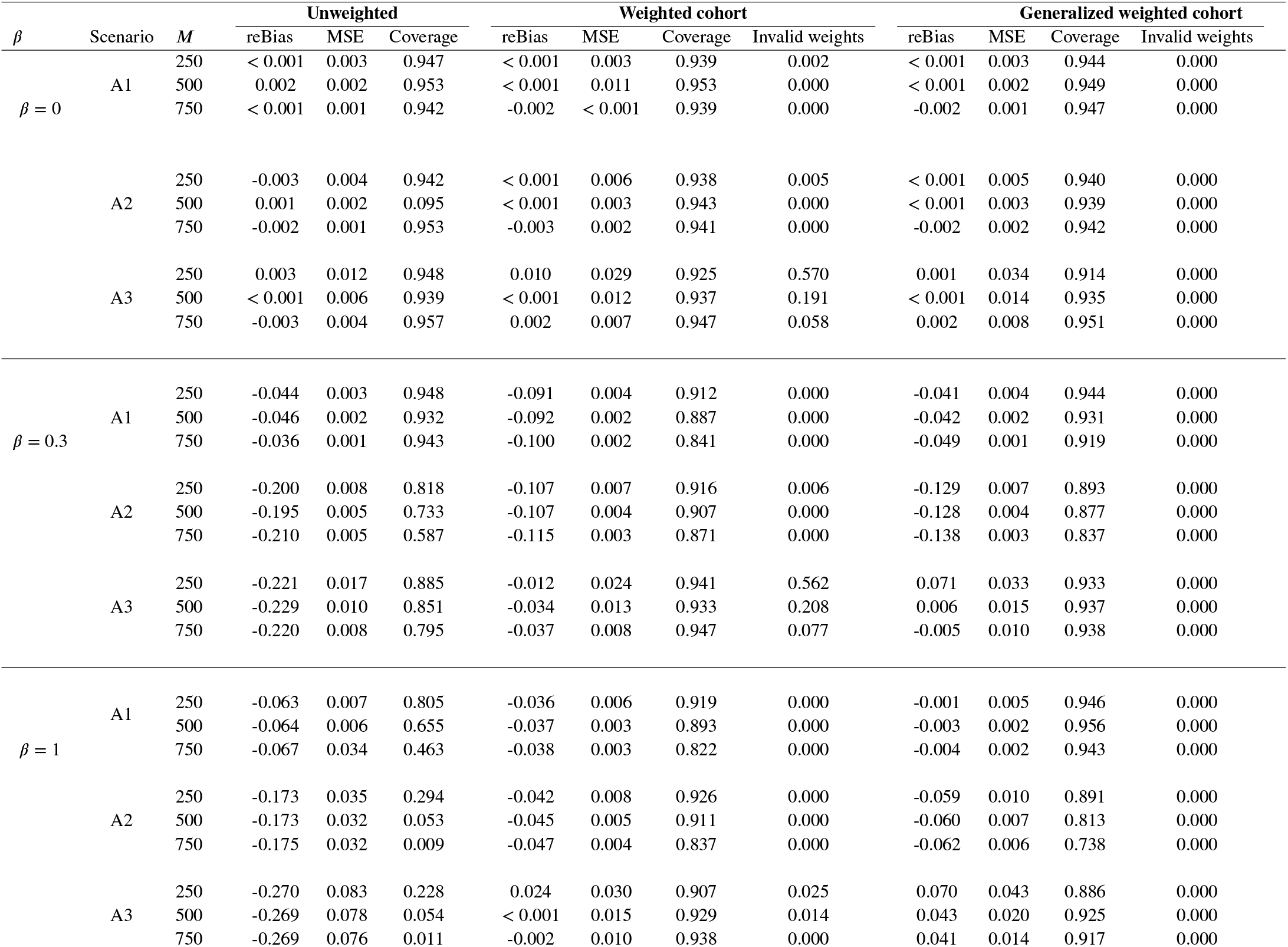
Simulation I. Relative bias (reBias), mean square error (MSE) and coverage probability (Coverage) for 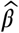 along 1000 trials. A1: mild level of ascertainment. A2: medium level of ascertainment A3: strong level of ascertainment. *M*: number of families.

### 3.2 Simulation setup II

In the previous simulation setting, we have assumed, as the proposed models in Section 2, that differences among individuals in terms of hazards can be fully accounted for by including covariates in the Cox proportional hazards model. However, when samples contain multiple members of the same family (often the case when applying the traditional weighted cohort approach as shown in Tabl S1), unmeasured hetereogeneity may arise since members of the same family often share common unmeasured characteristics such as genetic, social, dietary or other factors. In this second simulation setting, in order to introduce such unmeasured heterogeneity in the simulated data, we consider an extension of the data generation model specified in expression (14) by adding a latent (frailty) term, *U*, shared by all members of the same family:

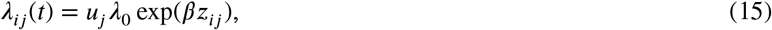

where *u*_*j*_ ∼ Γ(1, *θ*) is a latent term (frailty), shared by the *n*_*j*_ members of a given family *j*. The larger the value of the variance *θ*, the more family members are alike and the larger the difference between families, yielding larger unobserved family effects. We consider two different values of within-family correlation: ‘low’ (*θ*=0.1) and ‘large’ (*θ* = 1). Note that the latent frailty *U* and the covariate under investigation, *Z* are independent. We expect that as in the traditional unweighted Cox regression context (6), the presence of *U* introduces bias in the hazard ratio estimation, even when *U* is independent of the covariate of interest *Z*. However, we also expect that such a bias diminishes under the presence of outcome-dependent sampling.

The rest of the simulation settings were as in the previous Simulation setting I, except *M*, the number of families was fixed to 500 in this setting. For each scenario, weighted Cox models were estimated using the traditional and the generalized weighting scheme. Results obtained with the standard choices of using an unweighted Cox model or a shared gamma frailty model (unweighted) are also reported. Note that the application of the weighted approaches is not possible in the context of frailty models since the correct estimation of the weights would require knowing the true value of the frailty variance, which cannot be correctly estimated under outcome-dependent sampling.

### 3.3 Simulation results

#### 3.3.1 Simulation I

We first present the results obtained when data is generated under the assumption of fully observed heterogeneity. Both the level of outcome-dependent sampling and the covariate effect size determine the observed differences among the studied methods. If the covariate effect is strong (*β* = 1), the naive unweighted method is outperformed by the weighted approaches, even when the level of outcome-dependent selection is low (scenario A1). When the covariate effect is weaker (*β* = 0.3), and the level of ascertainment is medium (scenario A2) or high (scenario A3), both weighted cohort methods perform similarly and clearly outperform the naive, unweighted approach. Under a weak level of ascertainment (scenario A1), the new generalized weighted cohort method performs as well as the naive unweighted approach and they slightly outperform the traditional weighted cohort approach. Importantly, the traditional weighted approach is often not applicable when the assumed covariate effect is weak.

Negative weights are often obtained in this setting due to violation of condition (10). The same problem is observed in the null case scenario when assuming *β* = 0. The new generalized weighted cohort method does not suffer from this problem, yielding valid and satisfactory results in all studied scenarios.

#### 3.3.2 Simulation II

Table 2 shows the results assuming the presence of unobserved family-shared heterogeneity. When unmeasured within-family correlation is mild (*θ* = 0.1), we found similar results as in the previous simulation study: weighted methods perform similarly and provide better results than the unweighted model. Also, it is remarkable that weighted methods outperform the fit of a gamma shared frailty model which deals with shared unobserved heterogeneity but ignores outcome-dependent sampling.

**Table 2.** Simulation II. Relative bias (reBias), mean square error (MSE) and coverage probability (Coverage) for 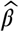 along 1000 trials. A1: mild level of ascertainment. A2: medium level of ascertainment A3: strong level of ascertainment. *M*: number of families. Data is generated according to a shared frailty model with frailty variance *θ*.

| $\theta$ | $\beta$ | Scenario | $M$ | Unweighted | | | Weighted cohort | | | | Generalized weighted cohort | | | | Shared frailty | | |
| --- | --- | --- | --- | --- | --- | --- | --- | --- | --- | --- | --- | --- | --- | --- | --- | --- | --- |
|  |  |  |  | reBias | MSE | Coverage | reBias | MSE | Coverage | Invalid weights | reBias | MSE | Coverage | Invalid weights | reBias | MSE | Coverage |
| $\theta = 0.1$ | $\beta = 0$ | A1 | 500 | 0.002 | 0.002 | 0.937 | 0.003 | 0.002 | 0.949 | 0.000 | 0.003 | 0.002 | 0.953 | 0.000 | < 0.001 | 0.002 | 0.939 |
|  |  | A2 | 500 | < 0.001 | 0.002 | 0.953 | 0.003 | 0.005 | 0.952 | 0.000 | 0.003 | 0.004 | 0.952 | 0.000 | < 0.001 | 0.002 | 0.954 |
|  |  | A3 | 500 | < 0.001 | 0.006 | 0.945 | -0.002 | 0.012 | 0.941 | 0.036 | < 0.001 | 0.015 | 0.936 | 0.000 | 0.002 | 0.006 | 0.942 |
| | $\beta = 0.3$ | A1 | 500 | -0.065 | 0.002 | 0.918 | -0.092 | 0.002 | 0.895 | 0.000 | -0.060 | 0.002 | 0.934 | 0.000 | -0.069 | 0.002 | 0.914 |
|  |  | A2 | 500 | -0.208 | 0.006 | 0.705 | -0.087 | 0.005 | 0.926 | 0.000 | -0.119 | 0.004 | 0.896 | 0.000 | -0.217 | 0.006 | 0.688 |
|  |  | A3 | 500 | -0.239 | 0.011 | 0.824 | -0.037 | 0.013 | 0.951 | 0.206 | -0.017 | 0.016 | 0.941 | 0.000 | -0.228 | 0.011 | 0.848 |
| | $\beta = 1$ | A1 | 500 | -0.094 | 0.010 | 0.363 | -0.059 | 0.006 | 0.766 | 0.000 | -0.032 | 0.003 | 0.895 | 0.000 | -0.092 | 0.010 | 0.394 |
|  |  | A2 | 500 | -0.195 | 0.040 | 0.027 | -0.057 | 0.007 | 0.865 | 0.000 | -0.080 | 0.010 | 0.734 | 0.000 | -0.194 | 0.049 | 0.030 |
|  |  | A3 | 500 | -0.278 | 0.083 | 0.045 | -0.012 | 0.017 | 0.921 | 0.000 | 0.017 | 0.022 | 0.912 | 0.000 | -0.278 | 0.083 | 0.063 |
| $\theta = 2$ | $\beta = 0$ | A1 | 500 | 0.001 | 0.002 | 0.945 | < 0.001 | 0.003 | 0.940 | 0.000 | 0.001 | 0.002 | 0.939 | 0.000 | < 0.001 | 0.006 | 0.954 |
|  |  | A2 | 500 | < 0.001 | 0.002 | 0.944 | -0.004 | 0.008 | 0.933 | 0.000 | -0.002 | 0.006 | 0.926 | 0.000 | -0.002 | 0.002 | 0.957 |
|  |  | A3 | 500 | 0.001 | 0.007 | 0.947 | -0.003 | 0.025 | 0.940 | 0.000 | -0.002 | 0.023 | 0.912 | 0.000 | -0.001 | 0.007 | 0.950 |
| | $\beta = 0.3$ | A1 | 500 | -0.238 | 0.007 | 0.613 | -0.130 | 0.005 | 0.885 | 0.000 | -0.164 | 0.006 | 0.854 | 0.000 | -0.131 | 0.004 | 0.830 |
|  |  | A2 | 500 | -0.279 | 0.009 | 0.567 | 0.070 | 0.008 | 0.939 | 0.070 | -0.043 | 0.006 | 0.940 | 0.000 | -0.231 | 0.007 | 0.667 |
|  |  | A3 | 500 | -0.310 | 0.015 | 0.793 | 0.076 | 0.044 | 0.879 | 0.480 | -0.051 | 0.039 | 0.899 | 0.000 | -0.323 | 0.016 | 0.756 |
| | $\beta = 1$ | A1 | 500 | -0.263 | 0.072 | 0.001 | -0.151 | 0.028 | 0.391 | 0.000 | -0.177 | 0.037 | 0.262 | 0.000 | -0.110 | 0.015 | 0.391 |
|  |  | A2 | 500 | -0.293 | 0.090 | 0.001 | 0.001 | 0.010 | 0.908 | 0.000 | -0.081 | 0.014 | 0.794 | 0.000 | -0.188 | 0.040 | 0.123 |
|  |  | A3 | 500 | -0.355 | 0.135 | 0.024 | -0.045 | 0.040 | 0.858 | 0.492 | -0.100 | 0.051 | 0.841 | 0.000 | -0.355 | 0.134 | 0.028 |

For strong within-family correlation (*θ* = 2) the performance of both weighted methods is, in general, good if the level of ascertainment is moderate or high (scenarios A2 and A3). If the level of ascertainment is mild (scenario A1), weighted methods would still outperform the traditional unweighted Cox approach but a shared frailty model seems a better choice in this setting. Bias is still noticeable with the shared frailty model, but of a smaller magnitude. Finally, the original weighted cohort also provided negative weights in this setting with unobserved family-shared heterogeneity, while the newly proposed generalized weighted cohort method proved to be more robust.

Overall, the simulation results indicate that the new generalized weighted cohort method is preferred over the original weighted cohort approach proposed by Antoniou et al. (2005). The original weighted cohort method performs, in general, well in presence of the combination of a strong covariate effect and strong outcome-dependent sampling, as expected. However, its applicability is restricted to certain scenarios and is not general enough. Inverse probability of selection weighted Cox models seem to still perform properly under the presence of mild unobserved family-shared heterogeneity, but they lead to biased results when the size of the frailty variance is large. Still, weighted methods seem to be preferred to the alternative approach of ignoring outcome-dependent sampling and fitting a shared frailty model if the level of outcome-dependent sampling is strong. If the level of ascertainment is mild, the results indicate a preference for the shared frailty model.

### 3.4 Software implementation

The generalized weighted cohort method developed in this work was implemented in the user-friendly R package ‘wcox’, which can be downloaded from https://github.com/vharntzen/wcox.

## 4 REAL DATA APPLICATIONS

We present two applications to illustrate the performance of the new generalized weighted cohort method compared to the traditional approaches on real data. In both applications, the goal is to assess the association between common susceptibility *loci* (gene locations on the chromosome) identified in Genome Wide Association Studies (GWAS) and cancer, using data collected through genetic testing in clinical genetics units. Specifically, the first application is devoted to study the association between a Single Nucleotide Polymorphism (SNP) and colorectal cancer (CRC) in carriers of a pathogenic variant in the *PMS2* gene while the second one focuses on the association of a 161 SNP-based polygenic risk score with breast cancer. The selection of both datasets was based on family history of cancer with oversampling of cancer cases with the aim of finding carriers of certain genetic variants. As a result, the sample used in the first application is composed of *PMS2* mutation carriers. In the second application, the sample is composed of high-risk families without BRCA1 or 2 mutations.

### 4.1 Application to colorectal cancer

In this application, we consider a sample of male carriers of the germline PMS2 mutation. Motivated by the previous promising findings reported by (7), we studied the association between the SNP rs1321311 and colorectal cancer in men. The sample consisted of 191 males belonging to 102 different families collected in eight Dutch clinical genetics centers between 2007 and 2016. Details on the selection criteria can be found in (8). The distribution of the family size was very skewed, the mean family size was 1.83 and most of the families (55 %) contributed with one single member (Figure 2, left panel). The last age of follow-up ranged between 25 and 88 years, but given that no events were observed after 75 years old, we censored observations at 75 years. The range of observed ages at CRC diagnosis varied between 25 and 75, and 58 events were observed. From the 191 studied individuals, 116 were homozygotes of the non-risk allele, 65 were heterozygotes and 10 were homozygotes of the risk allele.

**Figure 1.**
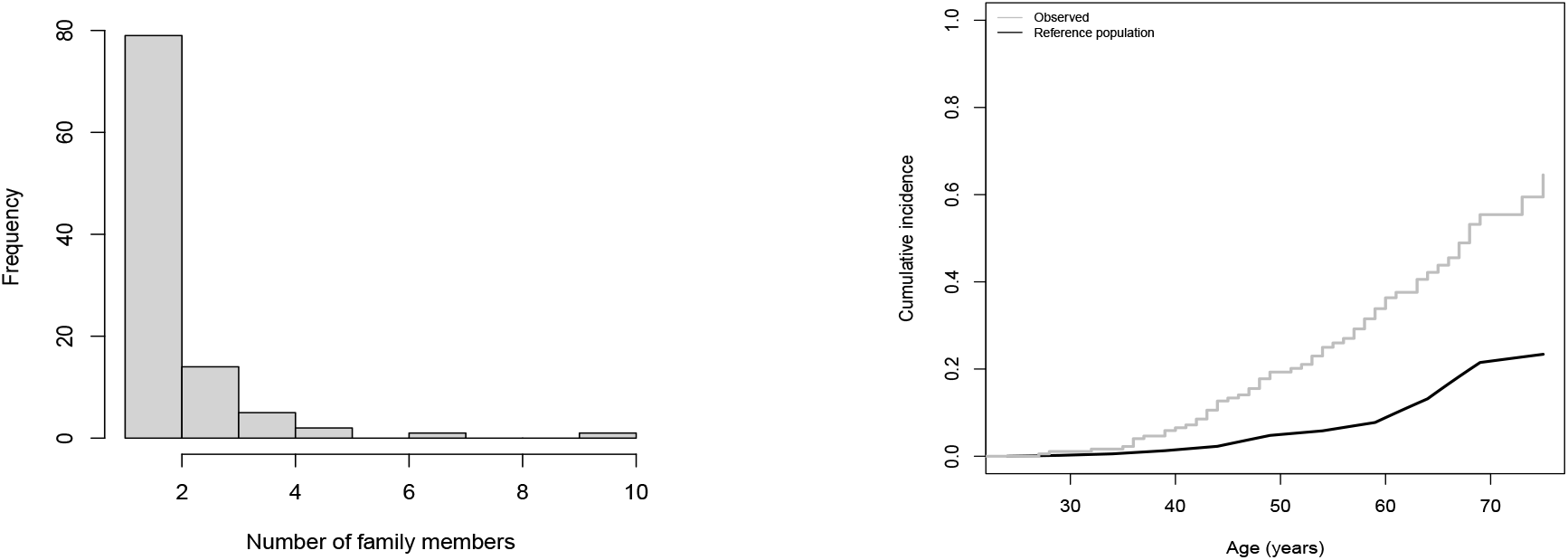
Application 1: Study of the association between SNP rs1321311 and CRC cancer in male carriers of a pathogenic variant in the gene *PMS2*. Left panel: Size of the families included in the sample. Right panel: Cumulative incidence of colorectal cancer at different ages. The gray line shows the observed risk in the sample. The black line reflects the expected cumulative colorectal cancer risk for the population of *PMS2* mutation carriers based on previous literature (8). Specifically, age-specific CRC incidence rates of *PMS2* mutation carriers are obtained multiplying the point estimates of the age-dependent hazard ratios as reported in Table 2 in (8) by the underlying population-based incidence rates of CRC for males in the Netherlands in 2011 according to the Netherlands Cancer Registry (NCR).

**Figure 2.**
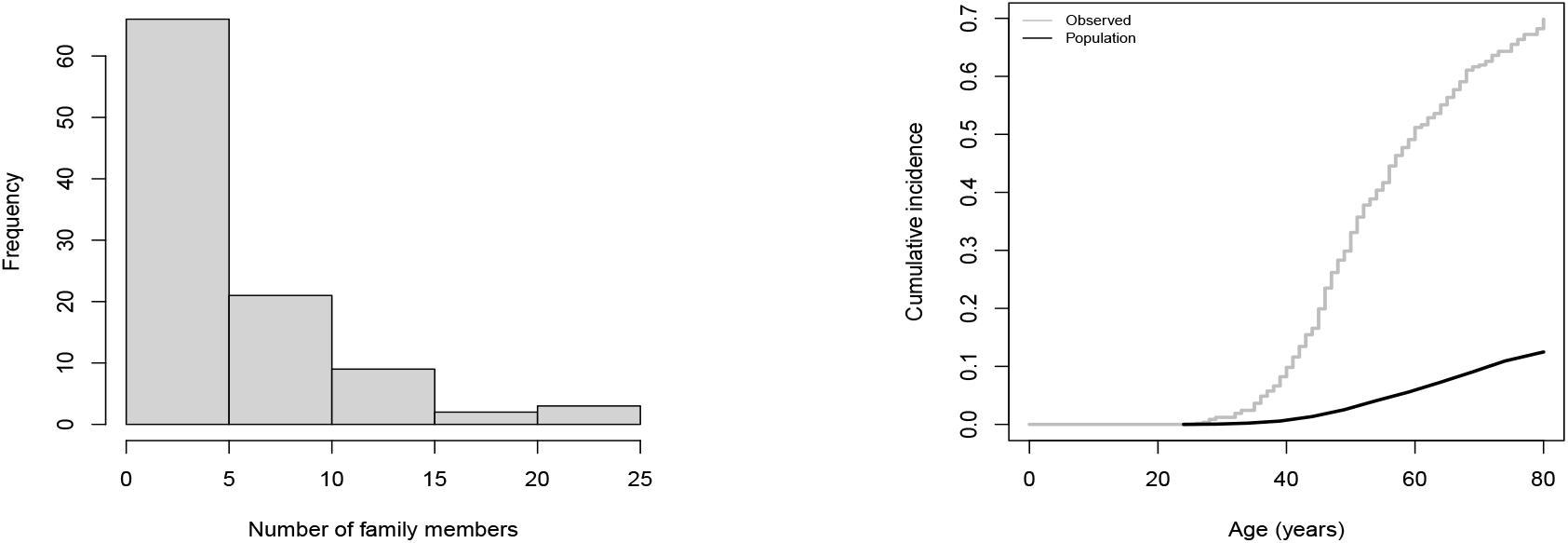
Application 2: Study of the association between a polygenic risk score and female breast cancer. Left panel: Size of the families included in the sample. Right panel: Cumulative incidence of breast cancer at different ages. The gray line shows the observed risk in the sample. The black line shows the population-based (the Netherlands, 2001 (9)) cumulative incidence used as reference in the weighted analyses.

Because of the limited size of the last category, we evaluated the effect of the indicator of being a carrier of the rs1321311 allele. We considered four different models: unweighted Cox regression, the state of the art weighted cohort method, our new method based on the new and more general weighting scheme and a shared gamma frailty model. The two studied weighted methods require the knowledge of incidence rates for CRC in carriers of pathogenic variants in *PMS2*. These were obtained by multiplying the population-based incidence rates of CRC in the Netherlands in 2011 (9) by the previously published ((8)) age-dependent hazard ratios of CRC for *PMS2* carriers. The choice of the year 2011 as the reference is justified because it is the middle point of the data collection period (2007-2016).

From the results reported in the bottom line of Table 3, it is observed that the new generalized weighted cohort method provides slightly larger estimated effects than the well-known (unweighted) Cox regression. In agreement to the result obtained with the unweighted method, the estimated association between the risk allele rs1321311 and CRC was statistically significant at the usual 5% level when using the new method. Importantly, the traditional weighted cohort approach could not be used because negative weights were obtained. Specifically, the oversampling of cases was not strong enough in the age group 65-70 years old and restriction (11) discussed in Section 2.1 was not met leading to negative weights for unaffected individuals in this age group. The shared frailty model provides the lower estimated covariate effect among the evaluated methods. This is probably due to the limited sizes of the family clusters and a small underlying unobserved heterogeneity. The estimated frailty variance was 0.15 with a broad confidence interval (0-1), indicating difficulties of the model to give reliable estimates of the level of unobserved heterogeneity. A likely major driving cause for this difficulty is the limited cluster size of this application since most of the families contribute a single individual to the analysis. As a consequence, the shared frailty approach is not recommended in this application and one would rather choose for the new generalised weighted cohort approach.

**Table 3.** Application to CRC in male carriers of PMS2. Estimated regression coefficients 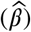 and corresponding 95% confidence intervals for the effect of the SNP rs1321311 for different Cox models. Case weights are calculated based on incidence rates of CRC for *PMS2* mutation carriers defined as the point estimates of the age-dependent hazard ratios reported in (7) multiplied by the population-based rates of CRC in Netherlands in 2011.

| Model | $\hat{\beta}$ (95% CI) |
| --- | --- |
| Unweighted | 0.723 (0.182-1.265) |
| Frailty | 0.671 (0.149-1.192) |
| Weighted cohort | - (*negative weights*) |
| Generalized weighted cohort | 0.771 (0.234-1.308) |

### 4.2 Application to breast cancer

In this application, the association between a PRS score and breast cancer was analyzed using a sample of 579 clinically ascertained women belonging to 101 families. On average, six women were included per family (mean family size = 5.73 and standard deviation = 4.66, Figure 2 right panel). The inclusion criterion was two-fold. Per family, one of the women should be tested negative for BRCA1 or BRCA2 pathogenic variants. This was a special feature of this sample and means that family aggregation and early-onset of cancer are not explained by pathogenic variants in these high-risk genes. Furthermore, breast cancer had to occur in at least three female family members or in two females if at least one had bilateral breast cancer before the age of 60. The families were selected between 1990 and 2012 by Clinical Genetic Services in four Dutch cities (Groningen, Leiden, Nijmegen and Rotterdam) and one Hungarian city (Budapest). Given the scarcity of events after 80 years of age (only one observed event at 90), we censored observations at age 80. The PRS was based on 161 SNPs weighted by previously published log-odds ratios (mostly based on population-based case-control studies). Detailed description of the calculation of this PRS can be found elsewhere (10). As before, to establish the association between the marker of interest, the PRS, and breast cancer, we considered four different models: the traditional unweighted Cox regression, the state of the art weighted cohort method to deal with outcome-dependent sampling, our new weighted method and a shared gamma frailty model. Population-based incidence rates of the Netherlands in 2001 (9) (mid point of the sample selection period) were used as external input to construct the weights.

From the results reported in Table 4, we observe that the new method provides a slightly smaller effect than the previously proposed weighted cohort approach and that both provided smaller effects than the unweighted Cox model. None of these three approaches reached statistical significance at the 5% level. In order to estimate the level of heterogeneity due to unmeasured within-family similarity, a shared frailty model was also fitted. The estimated frailty variance was 0.41, indicating that unobserved heterogeneity is not negligible in this application. This, together with the large size of the included families, is probably the reason why the shared frailty model seems to outperform the other methods. The estimated conditional hazard ratio using a shared frailty model is larger than the ones obtained using unweighted and weighted versions of the Cox model even if statistical significance at the 5% level is also not reached with this approach. According to our simulation results, we infer that the association between PRS and breast cancer is likely obscured by ignoring the strong unobserved heterogeneity and that the frailty approach is preferred in this application.

**Table 4.** Application to female breast cancer in non-BRCA1/2 families. Estimated regression coefficients 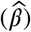 and corresponding 95% confidence intervals for the effect of polygenic risk score (PRS) for different Cox models.

| Model | $\hat{\beta}$ (95% CI) |
| --- | --- |
| Unweighted | 0.110 (-0.096, 0.317) |
| Frailty | 0.173 (-0.045, 0.390) |
| Weighted cohort | 0.079 (-0.226, 0.385) |
| Generalized weighted cohort | 0.062 (-0.261, 0.384) |

## 5 DISCUSSION

In this paper, we have revisited the analysis of outcome-dependently sampled survival data with weighted Cox regression using external data to construct inverse probability of selection weights. Our research is motivated by the interest in the effect of potential modifying factors on cancer risk using clinically ascertained data. Typically, those data sets are collected through ongoing genetic testing programs, where selection criteria lead to an overrepresentation of young cases and hence, the resulting samples are not representative of the target population of interest. We proposed a new weighting scheme that restores the expected number of events at each follow-up time using population-based hazard information. Our simulation study has shown that the new method can be applied to a broader set of realistic scenarios. Our real data applications support the same conclusion indicating the broader applicability of the new weighting scheme and it should be the preferred option to analyze data obtained under family-based outcome-dependent sampling when unobserved heterogeneity is negligible or mild.

A strength of the new weighting scheme is that it relies on fewer assumptions to provide valid, non-negative weights. The traditional weighted cohort (4) approach requires that a number of conditions are fulfilled, which hamper its applicability. Specifically, the original method is problematic if oversampling of cases is not observed in all age groups. In practice, although overall oversampling of events is expected, it does not necessarily hold for all age groups. Our new method overcomes this restriction and can be applied to a wider set of oversampling schemes, hence it can be regarded as a generalization of the traditional weighted cohort approach. This together with user-friendly implementation makes it an attractive analysis tool for applied researchers in the field.

Likewise the previously proposed weighted cohort method, our approach relies on a number of assumptions. First, a crucial assumption is the existence of a well-established external source of population-based incidence rates. Second, the sampling probabilities of observed individuals depend on the age at onset but they are assumed to be conditionally independent of the risk modifier under investigation. These two assumptions have been previously discussed in the context of the weighted cohort method (4, 5). An extra important assumption, but less often addressed, is fully observed heterogeneity. This means that if residual familiar aggregation is present, caution is required when interpreting the results. We have explicitly investigated this issue. Under the presence of unmeasured shared family factors affecting time to event, the estimation of a covariate effect is problematic, even if this covariate is independent of such unmeasured effects. The mere presence of such unobserved effects will affect the estimation of the hazard ratio of interest and careful interpretation of the results is required. Unobserved familiar heterogeneity induces a selection process based on the unobserved factors, so that the effect of the factor under investigation is attenuated with time, since the frailer individuals experience the event earlier in time. We have seen that also in this case, the use of weighted approaches seems advisable with respect to the naive unweighted approach. Also, if the number of available individuals per family is limited, i.e. cluster size is small, our new method might be the preferred option, outperforming a shared frailty model and the traditional weighted cohort approach. However, we would like to warn about the interpretation of the estimated effect and point out the systematic downwards bias of the regression coefficient in this setting. The extension of weighting approaches to the context of frailty models seems challenging. Since the estimated incidence rate in the sample depends on the correct estimation of the frailty variance, it would be necessary to know the value of the frailty variance to derive correct weights. However, the frailty variance is latent and hence we anticipate an identifiability problem in such an approach.

More sophisticated modeling, using a frailty model with explicit correction for ascertainment is possible but not straightforward and it is left as future research. It is noteworthy that such a complex approach will presumably require large clusters and sample sizes and hence our simpler approach based on borrowing information from a trustworthy external source will still be preferred in a number of relevant practical situations, such as our application to PMS2 carriers.

In conclusion, for performing regression analysis using survival data obtained under family-based outcome dependently sampling, specialized techniques are required to avoid bias and provide valid inference. We have proposed an accurate and conceptually simple method which generalizes and outperforms existing methods based on weighted Cox regression.

## ^0^ Abbreviations

GWAS: Genome Wide Association Studies
BRCA: breast (and ovarian) cancer associated (genes)
reBias: relative Bias
MSE: mean squared error
CRC: colorectal cancer
DAG: directed acyclic graph
SNP: Single Nucleotide Polymorphism (genetic variant)

## ACKNOWLEDGMENTS

The departments of Clinical Genetics and Human Genetics (LUMC, Leiden) are gratefully acknowledged for providing the breast cancer data set.

## SUPPORTING INFORMATION

The following supporting information is available as part of the online article:

**Figure S1.**
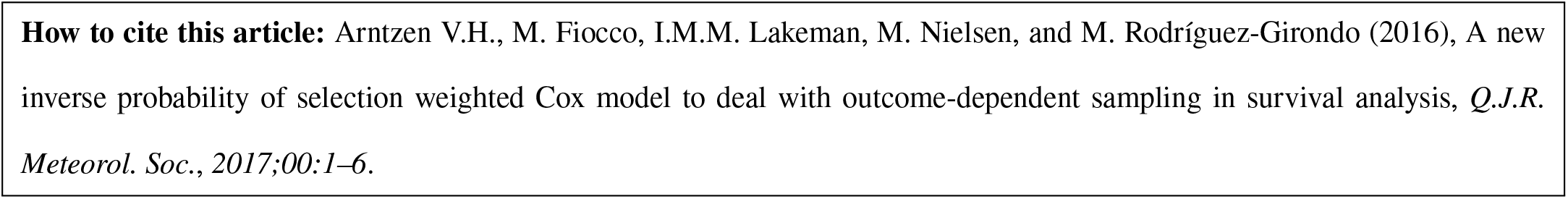
500 hPa geopotential anomalies for GC2C calculated against the ERA Interim reanalysis. The period is 1989–2008.

**How to cite this article:** Arntzen V.H., M. Fiocco, I.M.M. Lakeman, M. Nielsen, and M. Rodríguez-Girondo (2016), A new inverse probability of selection weighted Cox model to deal with outcome-dependent sampling in survival analysis, *Q*.*J*.*R. Meteorol. Soc*., *2017;00:1–6*.

## APPENDIX

## S1. LITERATURE REVIEW: USE OF WEIGHTED COHORT WITH FAMILY DATA

Among the 81 citations of Antoniou et al.’s 2005 paper (4), we found 51 papers that used a weighted cohort approach to obtain unbiased Hazard Ratios in a Cox model (list available upon request). The majority (62.7%, n = 32) were studies into breast and ovarian cancer risks. In 48 of the 51 papers applying the weighted cohort approach, the study sample certainly includes multiple members per family, but the exact number of families was only mentioned in 19 papers, shown in Table **??**. For each study, we calculated the average number of family members included. Note that this may fluctuate: one study (11) described the exact family composition of the sample, see Table **??** footnote 6. The median of the paper-specific, average family cluster sizes was

Generally, this data was collected at family canceror genetics clinics, where relatives of the index case (proband) were invited to be tested. Sometimes this was combined with ‘population-based’ recruitment (12, 13).

**Table S1:** Family information (when reported) in papers applying weighted cohort approach. This list is a subset of all (81) PubMed citations of the paper of *Antoniou et al*., 2005(4) on 06/02/202, with inclusion criteria 1) applying weighted Cox, 2) mentioning the number of families and sample size.

| *Authors* | *Year* | *Sample* | *Average number of relatives per family* | *(Cancer) research area* |
| --- | --- | --- | --- | --- |
| Borde et al.(14) | 2020 | 578 families; 760 carriers | 1.6 | Breast |
| Dashti et al.(15) | 2018 | 774 families; 2042 carriers | 2.6 | Ovarian |
| Ten Broeke et al.(7) | 2018 | 152 families; 521 samples | 3.4 | Colorectal |
| Kamiza et al. (16) | 2016 | 62 families; 260 carriers | 4.2 | Endometrial |
| Dashti et al. (17) | 2017 | 761 families; 1925 carriers | 2.5 | Colorectal |
| Chae et al. (12) | 2016 | 15049 families; 42489 participants<sup>(1)</sup> | 5.3 | Colorectal |
| Win et al. (18) | 2015 | 330 families; 1098 carriers | 3.3 | Breast and ovarian |
| Win et al. (19) | 2015 | 593 families; 854 individuals | 1.4 | General |
| Dashti et al. (20) | 2015 | 548 families; 1128 women | 2.1 | Breast |
| Ait Ouakrim et al. (13) | 2015 | 748 families; 1858 carriers<sup>(2)</sup> | 2.5 | Breast |
| Pooley et al. (21) | 2014 | 3134 families; 4822 included<sup>(3)</sup> | 1.5 | Breast |
| Killick et al. (22) | 2014 | 115 families; 158 included<sup>(4)</sup> | 1.4 | Breast |
| Win et al. (23) | 2013 | 315 families; 927 carriers | 2.9 | Breast |
| Pijpe et al. (24) | 2012 | 930 families; 1122 carriers<sup>(4)</sup> | 1.2 | Breast |
| Win et al. (25) | 2011 | 498 families provided 1324 carriers; | 2.7 (carriers); | Ovarian |
|  |  | 287 families provided 1219 non-carriers | 4.2 (non-carriers) |  |
| Win et al. (25) | 2011 | 286 families provided 601 carriers; | 2.1 (carriers); | Breast |
|  |  | 182 families provided 533 non-carriers | 2.9 (non-carriers) |  |
| Amos et al. (26) | 2010 | 93 families; 489 included (of which 45 married-in) | 5.3 | Ovarian |
| Antoniou et al. (27) | 2006 | 392 families; 810 carriers | 2.1 | Breast |
| Andrieu et al. (11) | 2006 | 1074 families; 1601 women | 1.5<sup>6</sup> | Breast |

(1) Includes some population-based recruitment for which average number of recruited relatives per family was 2.6, vs. 5.3 for clinic-based families.
(2) 25% population-based.
(3) Sampled non-carrying relatives only.
(4) Relatives functioned as controls, i.e. non-carriers.
(5) This concerns only one of the data sets in the paper.
(6) The relative occurrence of relatives per family was 71.1% for size 1, 17.9% for size 2, 6.2% for size 3, 2.8% for size 4 and 2.0% for size 5 up until 11.

## S2. POPULATION INCIDENCE RATES USED IN REAL DATA APPLICATIONS

**Table S2:** Table S2: Population-based age-specific incidence rates (in cases per 100.000) used for weight construction. For breast cancer, we used the registered data of The Netherlands in 2001 for women (9). For colorectal cancer, age-specific incidence rates were obtained by multiplying the population-based incidence rates of CRC in the Netherlands in 2011 (9) by the previously published ((8)) age-dependent hazard ratios of CRC for *PMS2* carriers. The choice of the year 2011 as the reference is justified because it is the middle point of the data collection period (2007-2016).

| Age group | Colorectal (men) | Breast (women) |
| --- | --- | --- |
| 25-30 | 40.56 | 9.09 |
| 30-34 | 69.39 | 33.61 |
| 35-39 | 146.60 | 72.41 |
| 40-44 | 204.38 | 156.17 |
| 45-49 | 518.28 | 241.48 |
| 50-54 | 221.10 | 323.56 |
| 55-59 | 411.68 | 310.83 |
| 60-64 | 1214.11 | 363.63 |
| 65-69 | 2016.21 | 388.41 |
| 70-74 | 405.12 | 415.42 |
| 75-79 | - | 293.22 |

